# Generalizing Michaelis–Menten Theory to Account for Substrate Heterogeneity

**DOI:** 10.1101/2025.08.21.671619

**Authors:** Poul Thrane, Johanne Gudmand-Høyer, Morten Andersen, Ulf Rørbæk Pedersen, William O. Hancock, Jeppe Kari

## Abstract

Enzymes are typically analyzed under the assumption of homogeneous substrates, yet many biological, biotechnological, and industrial reactions involve chemically or physically heterogeneous substrates. Here we present a theoretical framework showing that mixtures of non-identical substrates at steady state still follow the Michaelis–Menten (MM) rate law, yielding apparent parameters 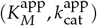 that depend on the mean and variance of the underlying energy distributions. This mathematical identity with the classical MM form can bias interpretation of fitted parameters and conceal mechanistic diversity. We show that variance in substrate energetics can shift 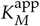 and 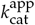 as strongly as changes in mean binding or activation energies, making substrate heterogeneity an independent but hidden design axis. Numerical simulations of 500 coupled Michaelis–Menten reactions—each representing a distinct substrate—validated the closed-form expressions for the apparent parameters. Analysis of 43220 curated BRENDA entries quantified the kinetic spread in enzymology, and our framework shows that realistic substrate heterogeneity gives comparable variability. Our framework extends the MM theory to heterogeneous substrates by adding a single, physically interpretable variance term to the classical equations. This enables inference of hidden heterogeneity from bulk fits and positions substrate pretreatment as a complementary optimization axis alongside conventional enzyme engineering for enzymes acting on heterogeneous substrates.

## Introduction

In recent years, single enzyme studies have advanced our understanding of enzyme kinetics by revealing varying catalytic efficiencies within a single enzyme population [1–5]. In parallel, recent advances in biocatalysis and metabolic engineering are shifting the focus of enzymology from mono-component enzyme reactions to multi-enzyme systems and network-level transformations involving complex substrates [6]. Unlike traditional enzyme kinetics, these composite systems introduce a new complexity characterized by the simultaneous presence of multiple substrates (or states of the substrate) and enzymes (or states of the enzyme) with a range of affinities, turnover numbers, and complex dynamics. It is critical to develop both models and methods to capture this kinetic heterogeneity, as ensemble-averaged measurements may conceal kinetic diversity and obscure the impact of heterogeneity in enzymology.

Most of the research on kinetic heterogeneity has focused on the enzyme itself [7, 8]. Numerous studies have elucidated the influence of conformational dynamics on enzyme binding [9] or activity [1–3, 10, 11], while other investigations have explored the “cocktail effect” involving multiple non-identical iso-enzymes within a complex mixture [12]. Less attention has been paid to the effect of substrate heterogeneity, although substrates can occupy long-lived conformational or physical states with different reactivity. For macromolecular substrates, this is well documented: protease digestion depends on the protein folding state, nuclease activity depends on nucleic-acid structure, and for insoluble polymers such as cellulose and PET, morphology and crystallinity strongly impact hydrolysis rates [13–16].

The technical and industrial applications of enzymes often involve enzymatic reactions in which numerous substrates coexist. For example, proteomics techniques, such as shotgun proteomics and quantitative proteomics, are based on enzymatic reactions, where a protease is incubated with a complex mixture of substrates (peptides or proteins) [17]. Other examples include interfacial enzymes that are important in the industrial application of enzymes [18, 19]. These enzymes typically act at the interface of an insoluble substrate, which can consist of multiple sub-sites with varying reactivities arising from differences in the morphology and crystallinity of the insoluble substrate [15, 16, 20–22].

Previous studies investigating the effect of substrate heterogeneity on enzyme kinetics have generally used a modeling approach in which each type of substrate in the mixture is individually parameterized [23–25]. Although this approach is feasible for systems with a few distinct substrates, it becomes impractical as the number of unique substrates increases, ultimately making it impossible to describe the system on an individual basis. Here, we examine the impact of substrate heterogeneity on steady-state enzyme kinetics in systems with multiple substrates. We begin by reviewing the classical Michaelis-Menten (MM) model and its energetic representation, and we generalize this view to enzyme reactions with multiple substrates using an energy distribution approach. The advantage of this approach is that substrate heterogeneity can be captured in a simple and scalable manner, using only one additional parameter (the variance of the distribution) compared to the classical MM model.

## Kinetic Model

### The Classical Case: Michaelis–Menten Kinetics with Homogeneous Substrate

The simplest and most popular model for the kinetics of an enzyme-catalyzed reaction is the Michaelis-Menten model.

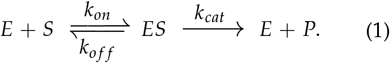

Here, a free enzyme (*E*) can bind to its substrate

(*S*) to form an enzyme-substrate complex (*ES*) governed by the second-order rate constant, *k*_*on*_. Once bound, the enzyme can escape the complex by dissociating back to *E* and *S* with a first order rate constant, *k*_*o f f*_, or turnover to form product (*P*) and *E* with the first order rate constant, *k*_*cat*_, called the turnover number. We denote the initial concentration of enzyme and substrate as *E*_0_ and *S*_0_, respectively. Applying the law of mass action, the law of mass conservation, *E*_0_ = [*E*] + [*ES*], Quasi-Steady-State Assumption (QSSA), *d*[*ES*]/*dt* = 0, and Reactant Stationary Assumption (RSA), [*S*] ≈*S*_0_, it is possible to derive the famous MM equation,

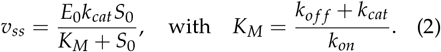

The MM equation connects the steady-state rate of product formation (*v*_*ss*_) of an enzyme-catalyzed reaction to two fundamental enzyme properties: stability, *K*_*M*_, and turnover number, *k*_*cat*_ of the enzyme-substrate intermediate.

An energetic representation of the MM reaction scheme is shown in Figure 1. The free energy diagram depicts the Gibbs free energy of binding (Δ*G*_*B*_) associated with the enzyme-substrate complex. When the dissociation rate of the *ES* complex to free *E* and *S* greatly exceeds the rate of product formation, i.e. *k*_*o f f*_ ≫ *k*_*cat*_, *K*_*M*_ can be treated as a dissociation constant (*K*_*D*_) for the *ES* complex [26] and Δ*G*_*B*_ is related to *K*_*M*_ by

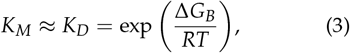

where *R* is the gas constant, and *T* is the temperature in Kelvin.

**Figure 1:**
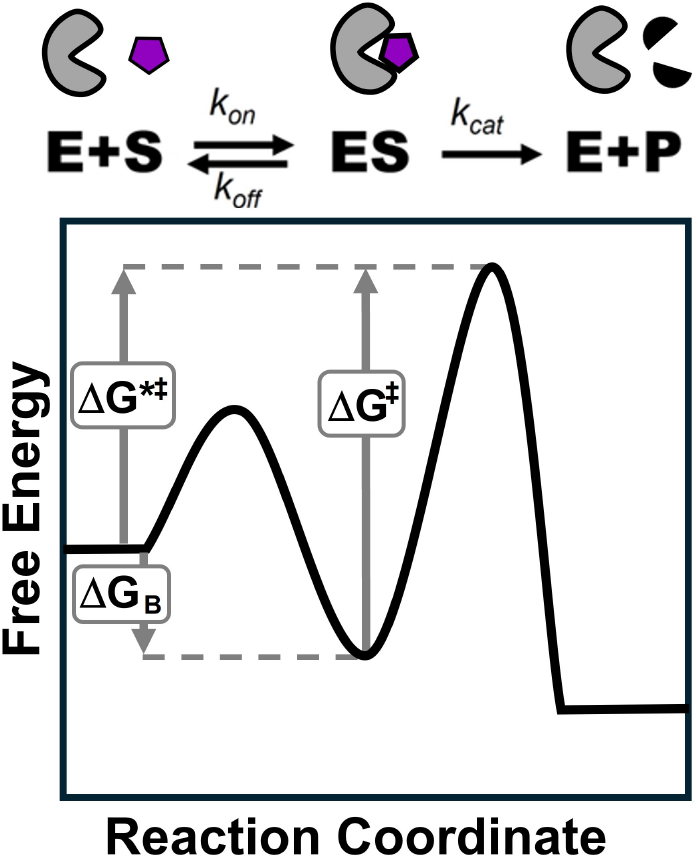
Energy diagram of an enzyme reaction following the Michaelis–Menten scheme. The diagram illustrates the variation in free energy along the reaction coordinate, from the free enzyme and substrate (E + S), through the enzyme–substrate complex (ES) and the transition state of the chemical step (ES^‡^), to the final state of free enzyme and product (E + P). The binding free energy (ΔG_B_), the activation free energy of the catalytic step (ΔG^‡^), and the overall energy barrier from the unbound state to the transition state (ΔG^*‡^ = ΔG_B_ + ΔG^‡^) are indicated in the diagram.

The Gibbs free energy of activation, Δ*G*^‡^, depicted in Figure 1, can be linked to the turnover number *k*_*cat*_, using the Eyring equation from transition state theory [27]

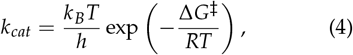

where *k*_*B*_ is the Boltzmann constant and *h* is Planck’s constant.

The ratio of the two MM parameters gives the specificity constant 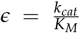. This parameter is important in enzymology, as it expresses the catalytic efficiency of the enzyme. It may be viewed as a second-order rate constant associated with the transition from the substrate to the product, with a barrier of Δ*G*^*‡^ = Δ*G*_*B*_ + Δ*G*^‡^ [26], as shown in Figure 1. From Eqs. (3) and (4), *ϵ* can be expressed as

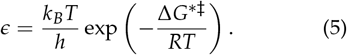

### A Generalized Michaelis–Menten Model with Heterogeneous Substrates

To generalize the MM model to include substrate heterogeneity, we extend Scheme (1) to *n* different substrates, as originally proposed by Schnell and Mendoza [23]. Specifically,

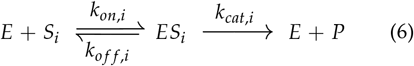

where *i* ∈ *{*1, 2, …, *n}*. In scheme (6), *S*_*i*_ is an individual substrate, *E* the free enzyme, *ES*_i_ the enzyme-substrate complex for each of the *n* substrates, and *P* the product. The initial enzyme and substrate concentration is *E*_0_ and *S*_0_ = ∑ *S*_*i*,0_, where *S*_*i*,0_ is the initial substrate concentration of *S*_*i*_. It is convenient to express *S*_*i*,0_ as a fraction of the total substrate, *S*_*i*,0_ = *S*_0_*α*_*i*_, where *α*_*i*_ denotes the initial mole fraction of *S*_*i*_.

Each reaction in scheme (6) is characterized by individual rate constants *k*_*on,i*_, *k*_*o f f*,*i*_, and *k*_*cat,i*_. This leads to *n* reactions coupled by a shared enzyme, where the overall reaction rate is given by the sum of the individual turnover rates for all enzyme-substrate complexes

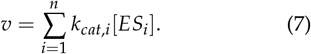

Using mass conservation for enzyme 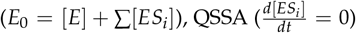, and RSA ([*S*_*i*_] ≈ *α*_*i*_*S*_0_), Eq. (7) simplifies at steady-state to (see SI for the full derivation)

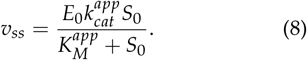

Where the apparent Michaelis-Menten parameters, 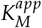 and 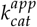, are defined as

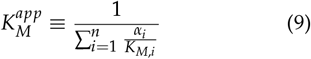

and

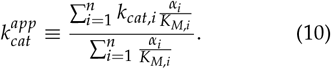

The apparent specificity constant can thus be defined as

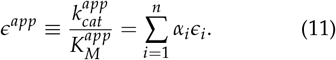

Remarkably, Eq. (8) is indistinguishable from the famous MM equation, Eq. (2), which is a new result to our knowledge. Thus, the generalized MM equation, Eq. (8), is applicable to enzyme-catalyzed reactions involving heterogeneous substrates, provided that the validity criteria for the generalized MM equation are met (see SI for details). This result emphasizes the remarkable robustness and versatility of the Michaelis-Menten formalism, which remains a cornerstone in enzymology [28]. However, Eq. (8) also underscores that the application of the MM equation on heterogeneous substrates could lead to severe misinterpretations of the MM parameters and hide the underlying complexity of the substrate, since the two equations have the exact same mathematical form. Thus, a small subpopulation of substrates with high affinity or high reactivity could disproportionately impact the observed 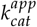 and 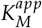.

The apparent MM parameters in Eqs. (9) and (10) initially appear complex; however, they exhibit a remarkable simplicity upon closer examination. Specifically, 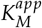 is the weighted harmonic mean (WHM) of the individual *K*_*M,i*_ (see SI), conceptually analogous to ensemble averaging in the binding-polynomial treatment of heterogeneous microstates [29]. Because the harmonic mean is dominated by the smallest terms, 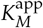 can be substantially smaller than the arithmetic mean when even a minor subpopulation has very low *K*_*M,i*_.

Similarly, defining the weights as 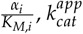 can be viewed as a weighted arithmetic mean (WAM) of the individual values of *k*_*cat*_, where the weight is the abundance to affinity ratio of each *S*_*i*_. Since the WHM and WAM are easy to compute for a continuous distribution, we will use this property of the derived apparent MM parameters in the following section and capture substrate heterogeneity using a free energy distribution approach.

### Energy representation of the generalized MM model

The energy diagram shown in Figure 1 is the classic textbook illustration of a simple homogeneous enzyme-catalyzed reaction. The energy diagram represents the lowest energy trajectory on an energy landscape, where high-energy barriers surround this specific pathway, ensuring that essentially all successful transitions occur along this dominant ‘highway’. This simplification works well when one pathway is by far the most likely. However, when multiple non-identical substrates are present, this picture breaks down. Figure 2 depicts this view of substrate heterogeneity. A narrow substrate ensemble behaves as if the substrate is homogeneous with similar affinity and reactivity, while a heterogeneous ensemble spans multiple substrate classes (five examples shown) with different kinetic properties.

**Figure 2:**
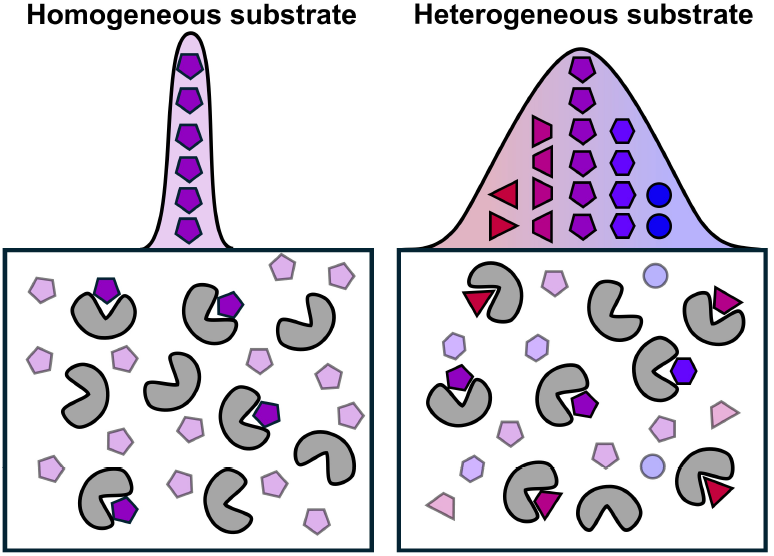
Conceptual illustration of an enzyme reaction with either homogeneous or heterogeneous substrates. In the homogeneous case, all substrates (purple pentagons) and enzymes (gray Pac-Mans) are similar, resulting in a narrow distribution of, e.g., binding affinities. In the heterogeneous case, the substrates and their kinetic properties vary more widely, leading to a broader distribution.

The generalized MM model in scheme (6) can be thought of as an extension of the single-pathway model to an energy landscape that contains *n* parallel reaction pathways. In this analogy, each of the *n* pathways represents the micro-path taken by a collection of substrates in the mixture that can be captured by the same rate constants (*k*_*on,i*_, *k*_*o f f*,*i*_, and *k*_*cat,i*_). This is visualized in Figure 3. In this representation, the substrate heterogeneity now manifests itself as a variance in the energy of the different states (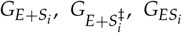, and 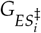) along the reaction coordinate. Thus, each unique substrate *S*_*i*_ will be associated with a reaction pathway given by the energy diagram *i*. Further, each substrate will be linked to a binding free energy 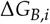 and an activation free energy 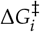, which can be converted to a *K*_*M,i*_ and *k*_*cat,i*_ using Eqs. (3) and (4).

**Figure 3:**
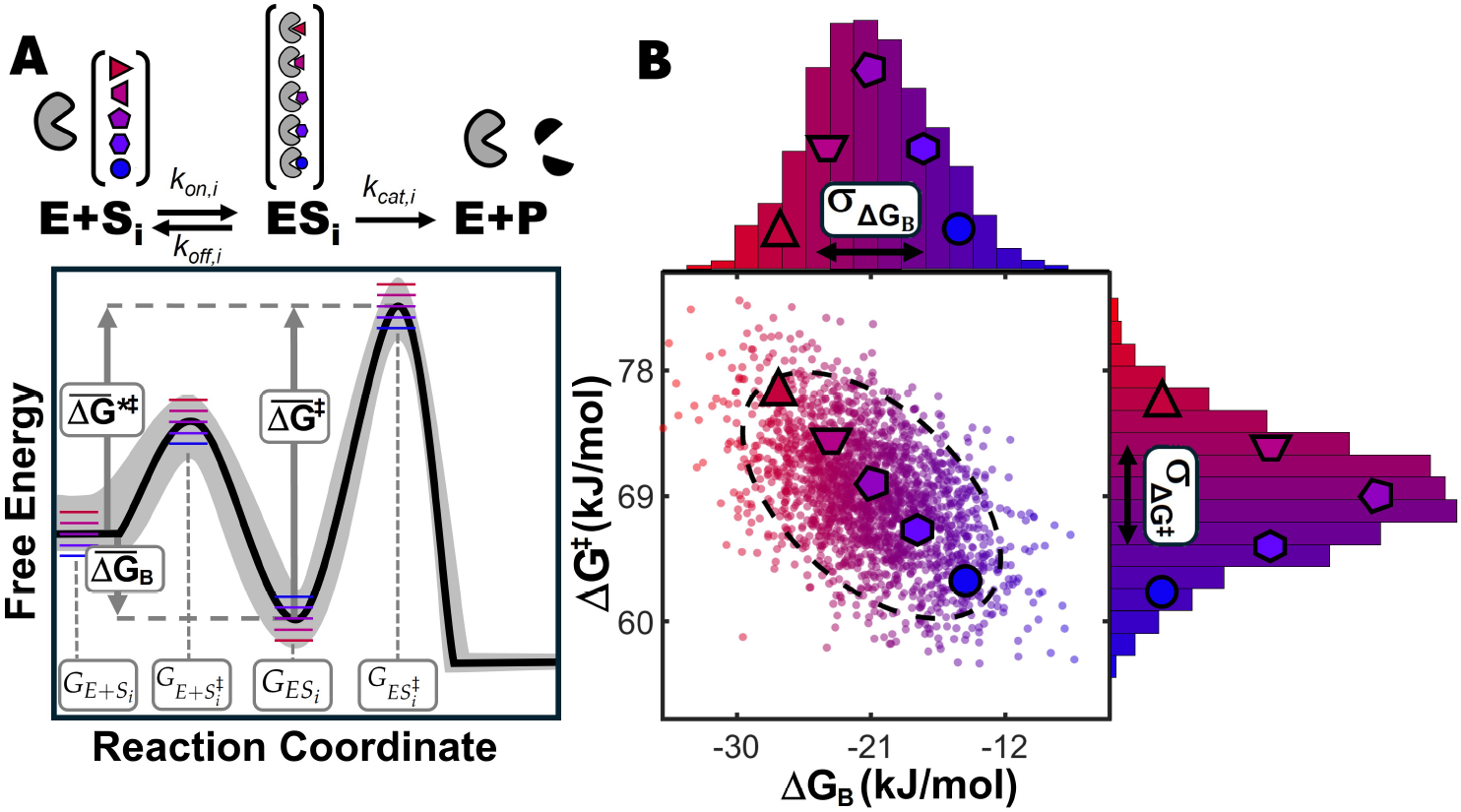
Generalized energy diagram for an enzyme-catalyzed reaction with heterogeneous substrates. A) Reaction scheme and corresponding energy diagram for the generalized MM model. Unlike the classical case shown in Figure 1, the free energy of all states and transition states (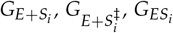, and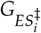) are normally dis-tributed across due to presence of multiple substrates. The gray band illustrates the variability in free energy around the mean energy profile (black curve), reflecting how different substrates populate the band. For illustration, five representative substrates with discrete energy levels are shown as colored lines in the diagram and as cartoon shapes above the reaction scheme. B) Scatter plot of the resulting free energy of binding (ΔG_B_) and activation (ΔG^‡^) for 500 randomly sampled energy diagrams based on the distributions shown in A. The five cartoon substrates from A are positioned according to their corresponding energy values. The purple pentagon substrate represents the mean binding and activation free energy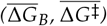. Standard deviations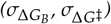 are indicated in the marginal histograms above the scatter plot. The dashed black ellipse denotes the 95% confidence region. Note ρ = −0.5), due to identical variance being applied to all states in the energy diagrams.

### Substrate heterogeneity as a continuous energy distribution

To examine the effect of substrate heterogeneity, we will assume that the free energy variations around the states (*G*_*E*+*S*_, *G*_*E*+*S*_‡, *G*_*ES*_, and *G*_*ES*_‡) follow independent normal distributions with means 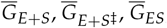, and 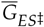 and variances 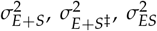 and 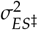. At the molecular level, this variation in energies may reflect differences in substrate conformation, configuration, or physical state. For example, a structured peptide may bind to a protease with affinity different from that of an unstructured peptide, and the crystalline regions of an insoluble polymeric substrate may impose energy barriers for an interfacial enzyme different from the amorphous regions. Modeling these differences as a continuous energy distribution allows us to simplify the expression of the apparent parameters from the statistical behavior of the ensemble. The assumption of normally distributed energies is reasonable since small and independent fluctuations around a mean follow the central limit theorem. The assumption is further supported by single-enzyme kinetic studies, in which the activation free energy of the enzyme population was found to be normally distributed [3, 11].

As shown in the Supporting Information, it is straightforward to derive an analytical expression for the apparent MM parameters when the free energies of all states in the energy diagram are normally distributed and time-independent. The analytical expressions for 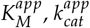 and *ϵ*^*app*^ are,

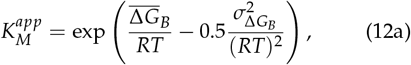

where

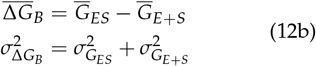

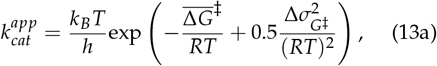

where

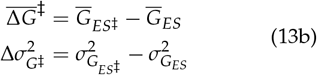

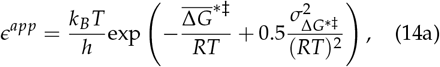

where

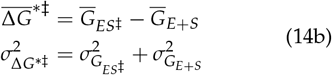

Eqs. (12a), and (13a) are of general interest as they connect the observable apparent MM constant 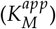 and turnover number 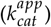 with the parameters of the underlying energy distribution. In comparison to the classical Eqs. (3), (4) and (5), the generalized MM parameters, Eqs. (12a), (13a) and (14b), only introduce one additional term related to the variance of the energy distributions that arise from the heterogeneity of the substrate (see Figure 2). In the limiting case where 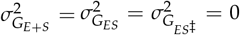, Eqs. (12a), (13a), and (14a) simplify to the classical expression for the MM parameters (Eqs. (3), (4) and (5)) in which the system can be treated as homogeneous with no or insignificant heterogeneity. Thus, the Gibbs free energy equation for an equilibrium constant (Eq. (3)) and the Eyring equation for a rate constant (Eq. (4)), may be viewed as a special case of Eqs. (12a), and (13a). In the next section, we will attempt to validate and apply Eqs. (12a), (13a), and (14a) to discuss the impact of substrate heterogeneity on steady-state kinetics. We note that although these equations were derived under the assumption of normally distributed free energies, the results also hold for other distributions (see SI).

## Results and Discussion

### Magnitude and Prevalence of Substrate-Driven Kinetic Variation

To investigate the prevalence and extent of substrate-driven kinetic heterogeneity, we first quantified the scale of kinetic diversity in enzymology and examined whether such levels of substrate-driven diversity are likely to exist. Figure 4A show the distributions of MichaelisMenten parameters extracted from the BRENDA database (accessed June 2025) [30]. After filtering the data (see Methods), the dataset contained *N* = 10590 values for *K*_*M*_, *N* = 14148 for *k*_*cat*_, and *N* = 18482 for *ϵ* (Figure 4A). Converting to energies via Eqs. (3), (4), and (5) yielded distributions well described by Gaussian fits with means 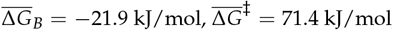, and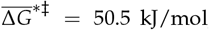, and standard deviations 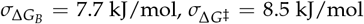, and 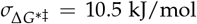(Figure 4B). These numbers help to quantify the magnitude of the kinetic diversity in enzymology. To put this span in relation to the current work, we also plotted literature values for two enzymes, *α*-chymotrypsin [31] and metallo-*β*-lactamase [32], assayed on different substrates (markers in Figure 4A–B). As seen from Figure 4 these individual MM parameters span nearly 90% of the kinetic diversity in the BRENDA database. Thus, variation among substrates recognized by one enzyme can give rise to significant substrate heterogeneity, motivating a model that explicitly accounts for substrate heterogeneity.

**Figure 4:**
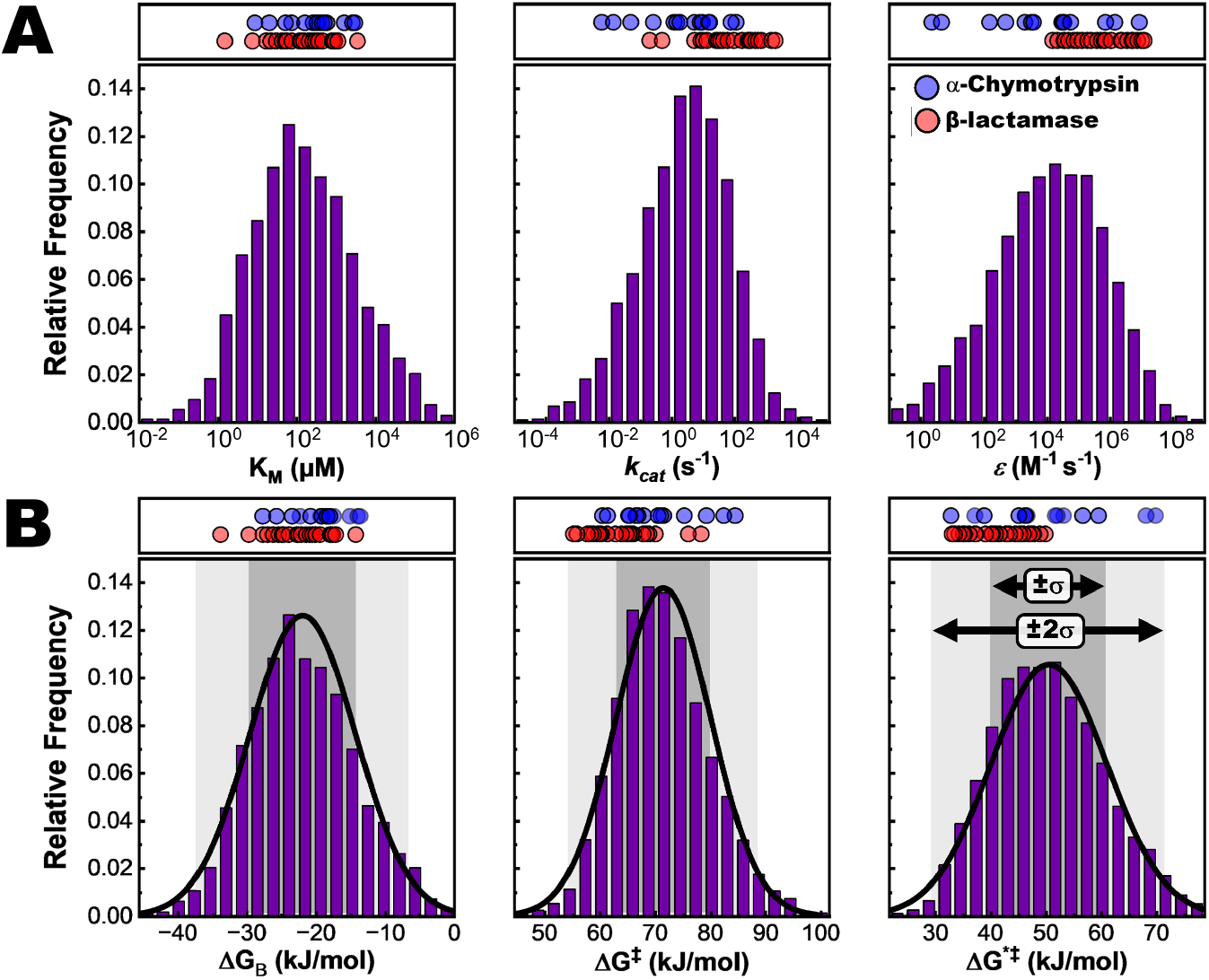
Distributions of enzyme kinetic parameters and their corresponding free energies. A) Michaelis–Menten parameters K_M_ (N = 10590), k_cat_ (N = 14148), and ϵ (N = 18482) extracted from the BRENDA database [30] (accessed June 2025). B) These parameters were converted to binding and activation free energies (ΔG_B_, ΔG^‡^, and ΔG^*‡^) using Eqs. (3), (4), and (5). The resulting energy distributions were fitted with Gaussian functions (black curves), yielding mean values of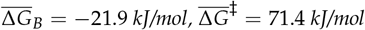, and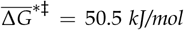, with standard deviations of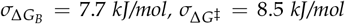, and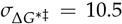 kJ/mol. Shaded regions indicate the 68% (±1σ) and 95% (±2σ) confidence intervals of the fitted distributions. Circle markers above the distributions in A and B represent literature-reported parameters and the corresponding calculated free energies for α-Chymotrypsin [31] (blue) and Metallo-β-lactamase [32] (red), each assayed on different substrates.

While the BRENDA analysis captures heterogeneity across chemically distinct substrates, single-molecule measurements have shown that substantial kinetic heterogeneity can arise even in apparently homogeneous systems. Mickert and Gorris [11] reported that *β*-galactosidase exhibits a normally distributed activation free energy with 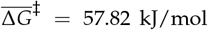 and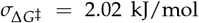. In that study, the heterogeneity was attributed to long-lived conformational states of the enzyme, but variations in the physical state of the substrate may likewise give rise to kinetic heterogeneity. The reported spread corresponds to a coefficient of variation of 3.5% 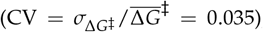, which is already sufficient in our framework to significantly shift the apparent parameters (Figure 5). Reactions on complex, heterogeneous substrates are therefore expected to introduce non-negligible heterogeneity that can systematically bias apparent kinetics, underscoring the need to explicitly incorporate substrate heterogeneity into kinetic measurements and analysis.

**Figure 5:**
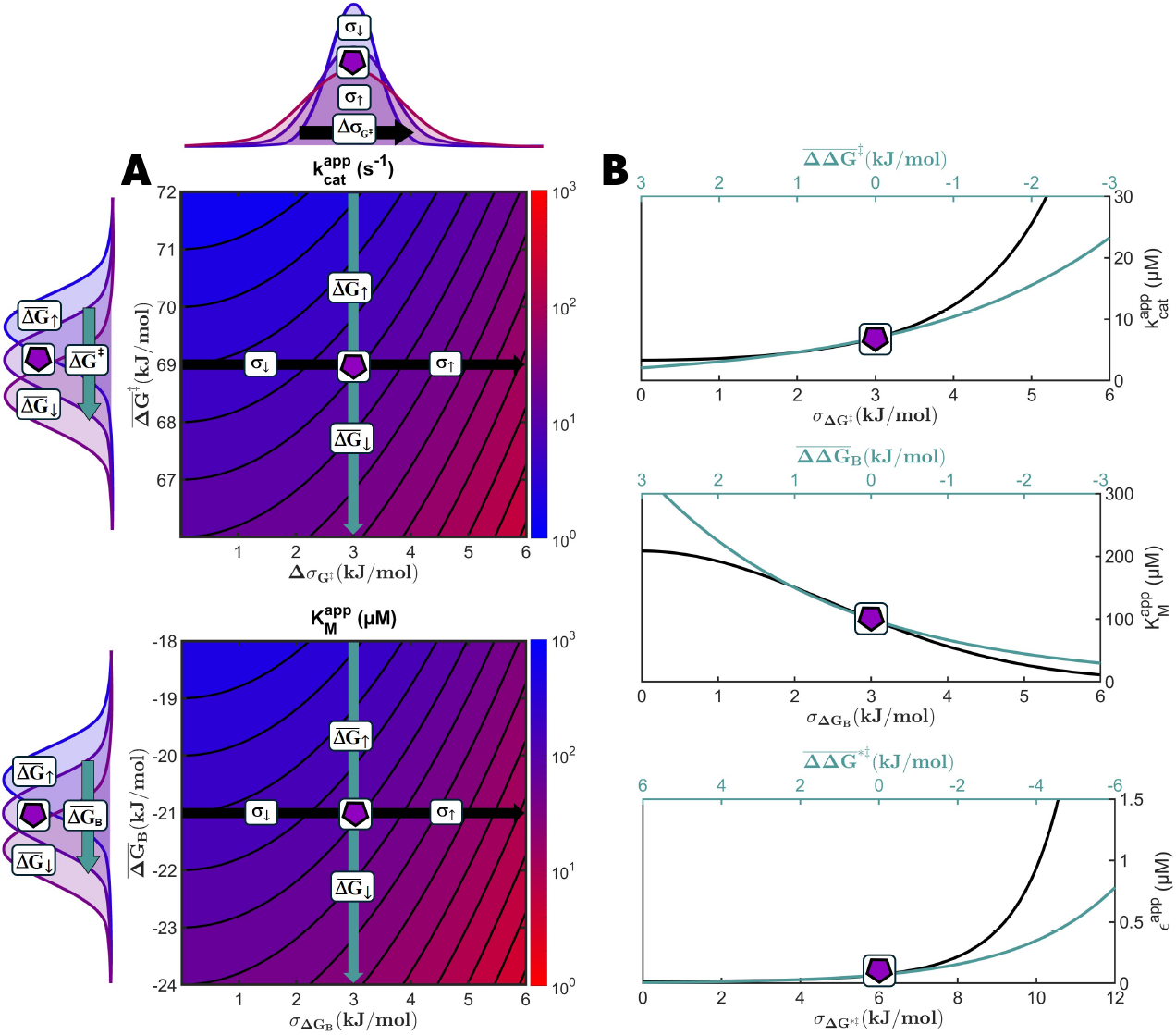
Impact on the apparent MM parameters by changes in the mean free energy (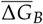 and 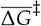) and standard deviation of the energy distribution (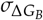 and 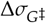;). A) Contour plots of Eqs. (12a) and (13a), illustrating how the apparent MM parameters change when the mean free energies and standard deviation are altered. B) Demonstration of the resulting apparent MM parameters when the mean or standard deviation is altered as given by the path of the teal and black arrows in A. The teal curve shows the effect of changing the mean free energy, while the black curve shows the effect of changing the standard deviation. Note that the energy span for both the mean energy (upper axis) and the standard deviation (lower axes) is the same. The purple pentagon in A and B illustrates the intersection of the vertical and horizontal arrows in A. The temperature in these plots was 25 ^°^ C and ΔG_B_ and ΔG^‡^ were assumed to be uncorrelated (ρ = 0). A scenario similar to the one shown in Figure 7A.

### Analytical predictions: substrate heterogeneity as a hidden design axis

To explore how substrate heterogeneity affects apparent kinetic parameters, we first evaluated the analytical expressions derived in Eqs. (12a) and (13a). The contour plots in Figure 5A makes the two independent parameters visible: vertical movements change the mean energies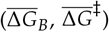, while horizontal movements change the heterogeneity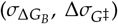. Whereas changes in the mean binding or activation energy predictably shift the apparent parameters, we also find that increasing heterogeneity alone can substantially enhance the apparent parameters (Figure 5B; black vs. teal curves).

Interestingly, the model predicts families of *iso-parameter contours* in the 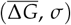plane: mean–variance pairs that leave 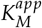 or 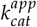un-change. Enzymes that lie on these iso-lines will appear kinetically identical although they may be mechanistically different. For instance, a fast and specialized enzyme (low 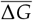 and *σ*) can fall on the same iso-line as a highly promiscuous and slow variant (high 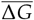and *σ*), making their bulk 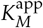 and 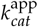indistinguishable. Screening for enzyme variants using ensemble-averaged kinetic measurements may not capture such trade-off in promiscuity and catalytic efficiency. Consistent with this notion, Sakuma et al. showed by single-molecule kinetics that certain *E. coli* alkaline phosphatase mutants exhibit increased functional-substate heterogeneity accompanied by enhanced promiscuous activities, an effect concealed in bulk measurements [33].

Another aspect of Figure 5A is the movement along the gradient perpendicular to an iso-line in the south-east direction (lower 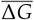 and higher *σ*) as this yields the largest changes in the apparent parameters. In this context, heterogene-ity can be viewed as an additional design axis that could complement traditional enzyme engineering of enzymes that act on heterogeneous substrates. Enzyme engineering may provide variants with higher catalytic efficiency (vertical movement in Figure 5A) and substrate pretreatment may broaden the substrate reactivity profile (horizontal movement in Figure 5A) to jointly increase the activity of the overall enzyme-catalyzed Interfacial enzymes such as amylases, cellulases, and PETases are examples of industrially relevant targets for such combined engineering strategy, since these enzymes have been shown to be highly sensitive to changes in the substrate morphology and crystallinity profile [15, 16, 20–22].

### Numerical simulation confirms analytical predictions

We validated the analytical predictions by numerical simulations of *n* = 500 parallel reactions parameterized by random draws from energy distributions of the different states in Scheme (6) (see Methods for details). For each heterogeneity setting (specified by 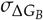 and 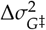) and cor-. relation regime (*ρ* = 0, −0.5, −1; Methods), we computed the complexation time course and ensemble-averaged steady-state rates.

As shown in Figure 6D–F, the substrate heterogeneity in the numerical simulations, was clearly visible in the pre-steady-state regime ([*ES*_*i*_](*t*)), whereas this complexity was hidden in the ensemble-averaged steady-state rates (*v*_*ss*_(*S*_0_)) shown in Figure 6G–I. As predicted, the overall steady-state rates retained the rectangular–hyperbolic form independently of the degree of heterogeneity. Hence, non-linear regression analysis gave perfect fits to Eq. (8) and recovered 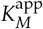 and 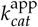 in quantitative agreement with the closed-form predictions of Eqs. (12a) and (13a) across heterogeneity levels and parameter correlations (Figure 7, lower panels). The dispersion around the apparent parameters derived from the numerical simulations increased with *σ* (error bars in Figure 7, lower panels) but was still in close agreement with the analytical predictions. The increase in noise was expected due to the finite sampling and sensitivity to high affinity and highly reactive substrates.

**Figure 6:**
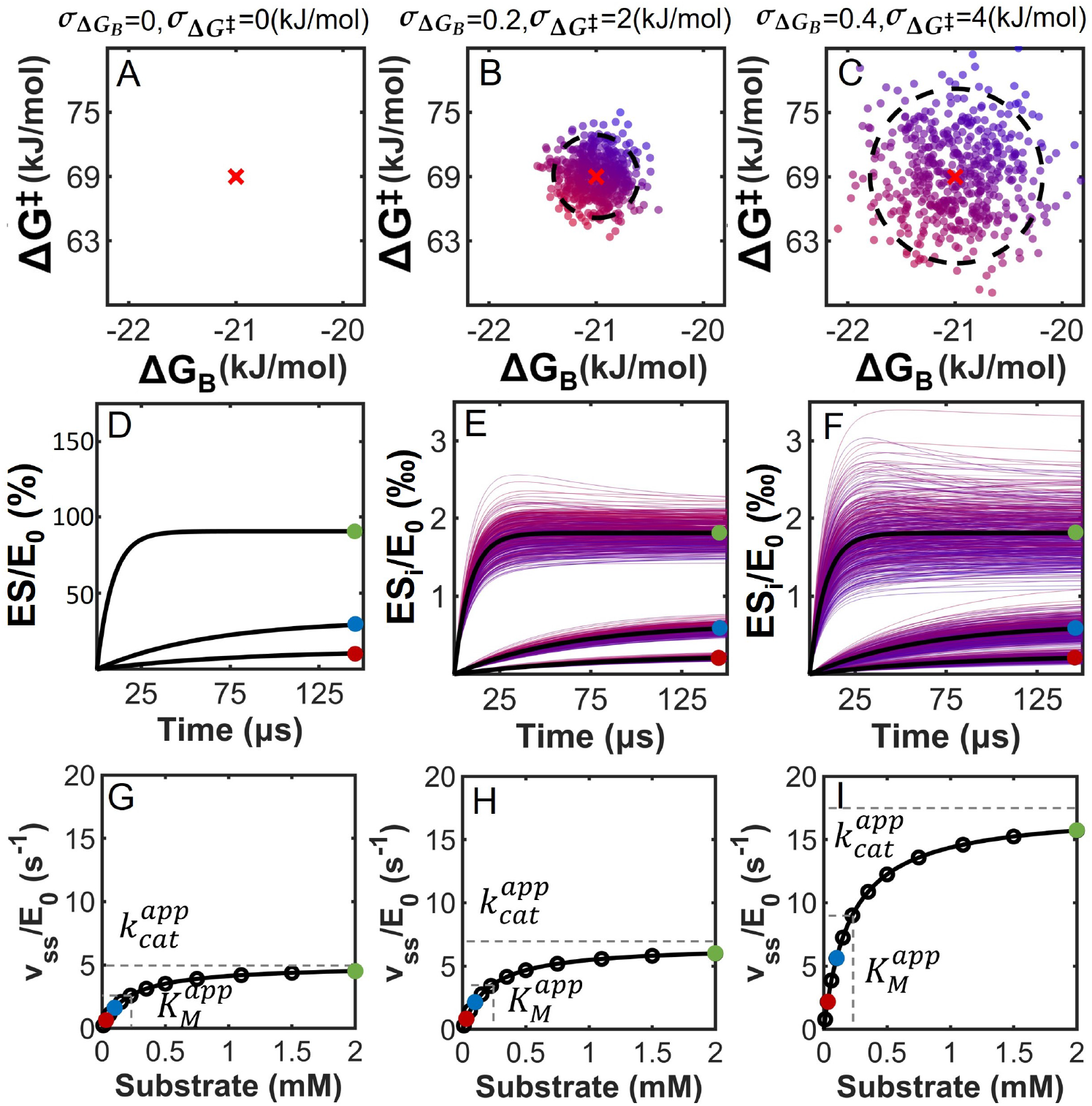
Effect of increasing kinetic heterogeneity on the enzyme-substrate complexation kinetics and steady-state rates. Upper panel (A–C): Scatter plots of binding (ΔG_B_) and activation (ΔG^‡^) free energies for 500 substrates (n = 500), sampled from independent normal distributions with increasing standard deviation (σ) from left to right. Red crosses indicate mean values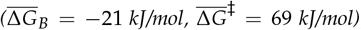; dashed ellipses show95% confidence intervals. Middle panel (D–F): Simulated complexation kinetics ([ES_i_]/E_0_) over time, based on the upper panel distributions. Each curve corresponds to one type of substrate, and the color gradient corresponds to the ΔG_B_ and ΔG^‡^ in the upper panel (A-C). Middle panel (D-F) includes three S_0_ values (0.03, 0.1, and 2 mM), marked with colored circles. These points match the substrate concentrations shown by the same colors in the lower panel. Lower panel (G–I): Apparent steady-state rates for increasing S_0_. Circles represent ensemble-averaged steady-state rates the numerical simulations and solid lines are the best fit to the generalized MM equation, Eq. (8), using non-linear regression analysis. In all simulations the total enzyme concentration was E_0_ = 100nM.

**Figure 7:**
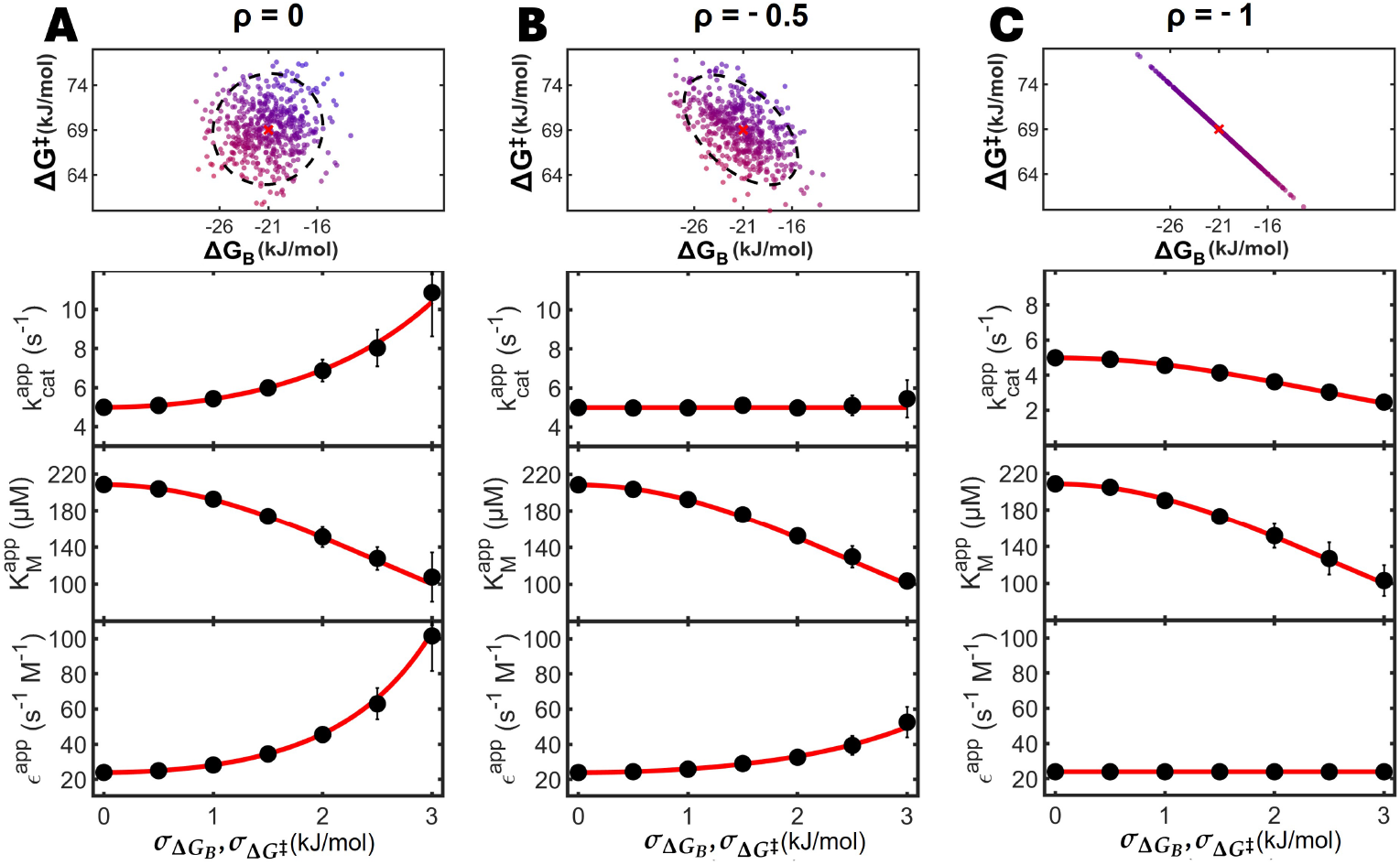
Comparison of apparent Michaelis–Menten parameters from analytical predictions and numerical simulations under varying substrate heterogeneity and energy correlation. The figure illustrates the close agreement between theory and simulation across different energy distributions and statistical couplings between binding and activation free energies. **Upper panels (A-C):** Scatter plots of bivariate normally distributed energies for 500 substrates, each with mean free energies 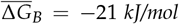 and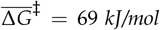, and standard deviations 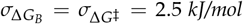ol. Correlation between ΔG_B_ and ΔG^‡^ increases from left to right: (A) uncorrelated (ρ = 0), (B) moderately anticorrelated (ρ = −0.5), and (C) perfectly anticorrelated (ρ = −1). Dashed circles/ellipses indicate 95% confidence intervals. Correlation is introduced by letting an increasing proportion of the total variance originate from the enzyme–substrate complex. **Lower panels (A–C):** Apparent Michaelis–Menten parameters (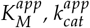, and ϵ^app^) obtained from simulations (black circles, error bars show 95% confidence intervals) and from analytical predictions (red curves) using Eqs. (12a), (13a) and (14a). The simulations were based on numerical integration of a coupled ODE system representing 500 parallel enzyme reactions (Scheme (6)), each parameterized by its own energy diagram drawn from the specified distributions.

### Parameter correlations and LFER depend on where in the energy diagram heterogeneity is manifested

We redistributed variance among *G*_*E*+*S*_, *G*_*ES*_, and 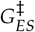to control the correlation between Δ*G*_*B*_ and Δ*G*^‡^ (Figure 7, upper panels). Three regimes emerge given by the Pearson correlation coefficient (*ρ*):

- *ρ* = 0 (variance in *E*+*S* and *ES*^‡^, none in *ES*): 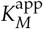 decrease and 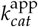 increase with hetero-geneity.
- *ρ* = −0.5 (variance shared equally across states); 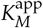decrease with heterogeneity, whereas 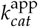 is unchanged.
- *ρ* = −1 (all variance assigned to the *ES* state): 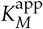and 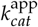decrease proportionally with increasing variance, leaving *ϵ*_app_ invariant.

Analytically, this follows directly from Eqs. (12a)–(13a): 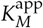depends only on the variance of Δ*G*_*B*_ (hence independent of *ρ*), whereas 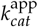 depends on 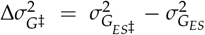 and is therefore sensitive to which state carries the heterogeneity. Concentrating variance in *ES* enforces an anticorrelation between Δ*G*_*B*_ and Δ*G*^‡^ (*ρ* = −1), yielding a constant *ϵ*^app^ as observed (Figure 7, lower panels).

Correlation between Δ*G*_*B*_ and Δ*G*^‡^ echo classic *linear free-energy relationships* (LFERs). When the substrate heterogeneity is primarily realized in the enzyme-substrate complex ensemble, shifts in binding and activation energies become linearly correlated. LFERs are well known in mechanistic organic chemistry and heterogeneous catalysis [34–37], but have seen limited application in biocatalysis [38]. Seminal work by Fersht et al. has shown how LFERs can provide mechanistic insight into the transition state of enzymes [39–42]. More recent studies have applied LFERs and the related Sabatier principle to rationalize the activity of interfacial enzymes working on insoluble substrates [43–45]. However, this body of work has traditionally focused on the enzyme rather than the substrate. Our results show that substrate heterogeneity alone can cause LFER between Δ*G*_*B*_ and Δ*G*^‡^ if the heterogeneity primarily manifest itself in the enzyme-substrate complex.

### Implications for measurement

Because Eq. (8) is mathematically indistinguishable from the classical Michaelis–Menten form, bulk steady-state assays alone cannot reveal substrate heterogeneity: mixtures with different underlying mechanisms may yield identical 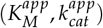.Our simulations show that heterogeneity is visible in the pre-steady-state regime but hidden once steady state is reached, underscoring that standard fits to Eq. (8) provide no diagnostic power on their own.

Three practical points follow:

- **Pre-steady-state or extended progress curves**. Deviations from single-exponential complexation kinetics, as seen in our simulations, are potential indicators of hidden heterogeneity. Ensemble steady-state rates alone are not sufficient.
- **Validity criteria**. In the Supporting Information we derived the condition 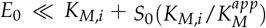which need to be fourfilled for all substrates in the mixture. This requirement is stricter than the homogeneous case (*E*_0_ ≪ *K*_*M*_ + *S*_0_), which may be exploited experimentally to test for substrate heterogeneity.
- **Reporting**. When heterogeneity is suspected, fitted 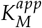 and 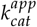 should be interpreted as ensemble parameters rather than intrinsic constants, and ideally reported together with an assessment of possible variance contributions.

Steady-state kinetics is essential in enzymology as many enzymatic reactions operate in this regime. Our framework directly connects substrate heterogeneity to the apparent steady-state parameters and show it impact on these parameters. However, bulk steady-state assays alone cannot determine whether a system is homogeneous or heterogeneous. To establish the presence and nature of heterogeneity, steady-state measurements should therefore be complemented with approaches that probe outside the steady-state regime, such as pre- or post-steady-state progress curves or single-molecule assays.

## Conclusion

We generalized Michaelis–Menten kinetics to mixtures of non-identical substrates by representing heterogeneity as distributions on the underlying energy landscape. Remarkably, heterogeneous and homogeneous systems are mathematically indistinguishable at steady state, as both yield the rectangular–hyperbolic MM form. However, in the heterogeneous case the apparent parameters reflect both the mean and variance of the underlying energy distributions. This identity explains why bulk fits can conceal mechanistic diversity and bias parameter interpretation.

Our analysis shows that variance alone can shift 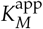 and 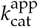 as strongly as changes in mean energetics, producing iso-parameter contours that render mechanistically distinct systems indistinguishable. When variance resides in the enzyme-substrate ensemble, binding and activation ener-gies become correlated, yielding LFER-like behavior. Numerical simulations and analysis of 43220 curated BRENDA entries indicate that realistic substrate heterogeneity is sufficient to reproduce database-scale dispersion. Because the framework is MM-compatible and introduces only one physically interpretable parameter (variance), it can be adopted immediately to improve mechanistic inference, inform experimental diagnostics, and support co-optimization strategies in which enzyme engineering and substrate pretreatment act as complementary design axes for enzymes operating on heterogeneous substrates.

## Methods

### BRENDA Database Analysis

Michaelis–Menten parameters (*k*_*cat*_, *K*_*M*_, and *ϵ*) were extracted from the BRENDA enzyme database (release 2025.1, accessed June 2025) [30]. To minimize temperature-related variability, only entries with reported assay temperatures between 20 and 30 ^*°*^C were retained, yielding *N* = 14148 for *k*_*cat*_, *N* = 10590 for *K*_*M*_, and *N* = 18482 for *ϵ*. The parameter distributions of the curated dataset were the same as the entire dataset (*k*_*cat*_, *N* = 32838, *K*_*M*_, *N* = 25484 and *ϵ, N* = 40376) and thus were considered representative of the parameter distribution of the full dataset. No additional filtering was applied with respect to organism, pH, buffer, or substrate. The curated *K*_*M*_, *k*_*cat*_ and *ϵ* values were converted to binding (Δ*G*_*B*_) and activation free energies(Δ*G*^‡^, and Δ*G*^*‡^) using Eqs. (3), (4), and (5) and each entry’s reported assay temperature. The resulting distributions were fitted to normal functions to obtain mean energies (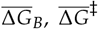and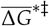) and standard deviations (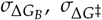and 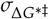). These fitted values were used to parameterize the simulations described below. Literature-reported *k*_*cat*_ and *K*_*M*_ values for *α*-Chymotrypsin [31] and Metallo-*β*-lactamase [32], assayed on multiple different substrates were overlaid on the BRENDA-derived distributions to highlight the kinetic variation that different substrates can give rise to.

### Numerical simulations

We simulated a system of *n* = 500 parallel enzyme–substrate reactions (Scheme (6)) with equal mole fraction (*α*_*i*_ = 1/*n*). Each reaction was assigned a unique energy diagram for the states 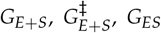 and 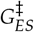. Energies were sam-pled from independent normal distributions with means: 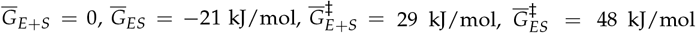, giving 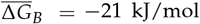 and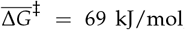, consistent with means from the BRENDA analysis and typical rate constant for enzymes (*k*_*on*_ ≈5 *×* 10^7^ M^−1^s^−1^, *k*_*o f f*_ ≈10^4^ s^−1^, *k*_*cat*_ ≈5 s^−1^) [26, 46, 47]. Microscopic rate constants (*k*_*on,i*_, *k*_*o f f*,*i*_, *k*_*cat,i*_) were obtained from the sampled energies using Eq. (4), yielding a coupled ODE system of 1002 equations. This system was solved in MATLAB (R2024b) using ode15s with *E*_0_ = 0.1 *µ*M and *S*_0_ ranging from 1 *µ*M to 1 M. The steady-state rate was computed using Eq. (7) and the apparent MM parameters where derived from fit to the generalized Michaelis–Menten equation (Eq. (8)). Each condition was repeated 10 times with independent random draws to estimate 95% confidence intervals. All simulations satisfied RSA and QSSA validity criteria for heterogeneous systems [23] (see SI for further details).

We defined the degree of substrate heterogeneity by the variances 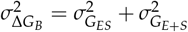 and 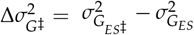 (Eq. (12b) and (13b)). Correlation between Δ*G*_*B*_ and Δ*G*^‡^ was imposed by systematically redistributing the total variance among energy states *G*_*E*+*S*_, *G*_*ES*_ and 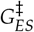. We examined three correlation regimes given by the Pearson correlation coefficient. *ρ* = 0: Total variance was distributed between *G*_*E*+*S*_ and 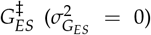. *ρ* = −0, 5: Total variance shared equally across states 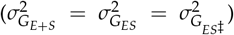. *ρ* = −1: Total variance assigned to 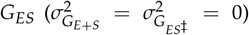. Examples of these distributions are shown in Figure 7A–C.

## Supporting Information

### Derivation of the generalized MM equation

We consider the system of coupled ODEs derived by applying the law of mass action on Scheme (6) (main article), given in compact form by

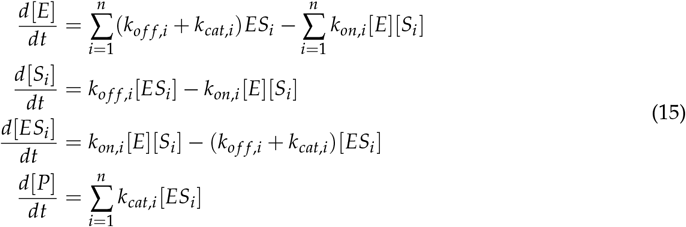

where *i* ∈ {1, 2, …, *n*}, [*E*] is the free enzyme concentration, [*S*_*i*_] is the concentration of the i’th substrate, [*ES*_*i*_] is the concentration of the i’th enzyme-substrate complex, and *k*_*on,i*_, *k*_*o f f*,*i*_ and *k*_*cat,i*_ are the local rate constants for the i’th reaction. Unfolding this yields a system of 2*n* + 2 coupled ODEs.

Applying the quasi-steady-state assumption 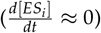 for all enzyme-substrate complexes, yields

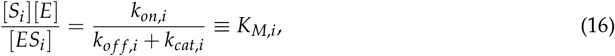

where *K*_*M,i*_ is the Michaelis-Menten constant for substrate *i*. The overall rate of the reaction is given by the sum of reaction rates for all individual substrates,

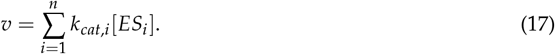

Isolating [*ES*_*i*_] in Eq. (16), and substituting into the rate expression above we get an expression for the overall steady-state rate (*v*_*ss*_)

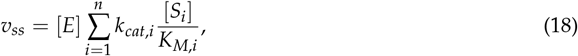

Since the enzyme is conserved,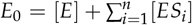, we can express the free enzyme concentration at steady-state as

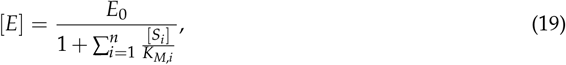

and combining Eqs. (18) and (19) gives

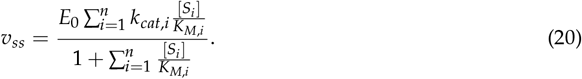

Assuming that the reactant stationary assumption (RSA) holds for all substrates ([*S*_*i*_] ≈ *α*_*i*_*S*_0_) this equation reduces to

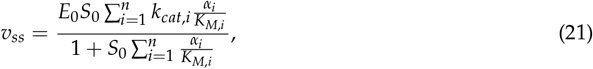

which can be rewritten as Eq. (8) in the main article, by defining the apparent kinetic parameters, 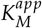 and 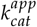, as in Eqs. (9) and (10). In the next section (Validity of the generalized MM equation) we show under which conditions the QSSA and RSA are valid and hence under which conditions the generalized MM equation can be applied.

### Validity of the generalized MM equation

To investigate under which conditions the generalized MM equation is valid, we start by looking at the valid timescale for QSSA. Schnell and Mendoza developed an expression for the short and long timescales for QSSA in a system with multiple substrates [23]. Adapting our notation to their expression gives

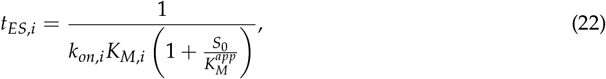

and

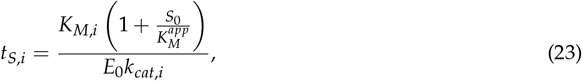

where *t*_*ES,i*_ is the (short) timescale for substrate *S*_*i*_ to reach steady-state, and *t*_*S,i*_ is the (long) timescale for the steady-state to end for substrate *S*_*i*_. These timescales were compared to numerical simulations which where setup as described in the main article. Results from these simulations are shown in in Figure 8D-I, and show that these timescales are indeed applicable.

**Figure 8:**
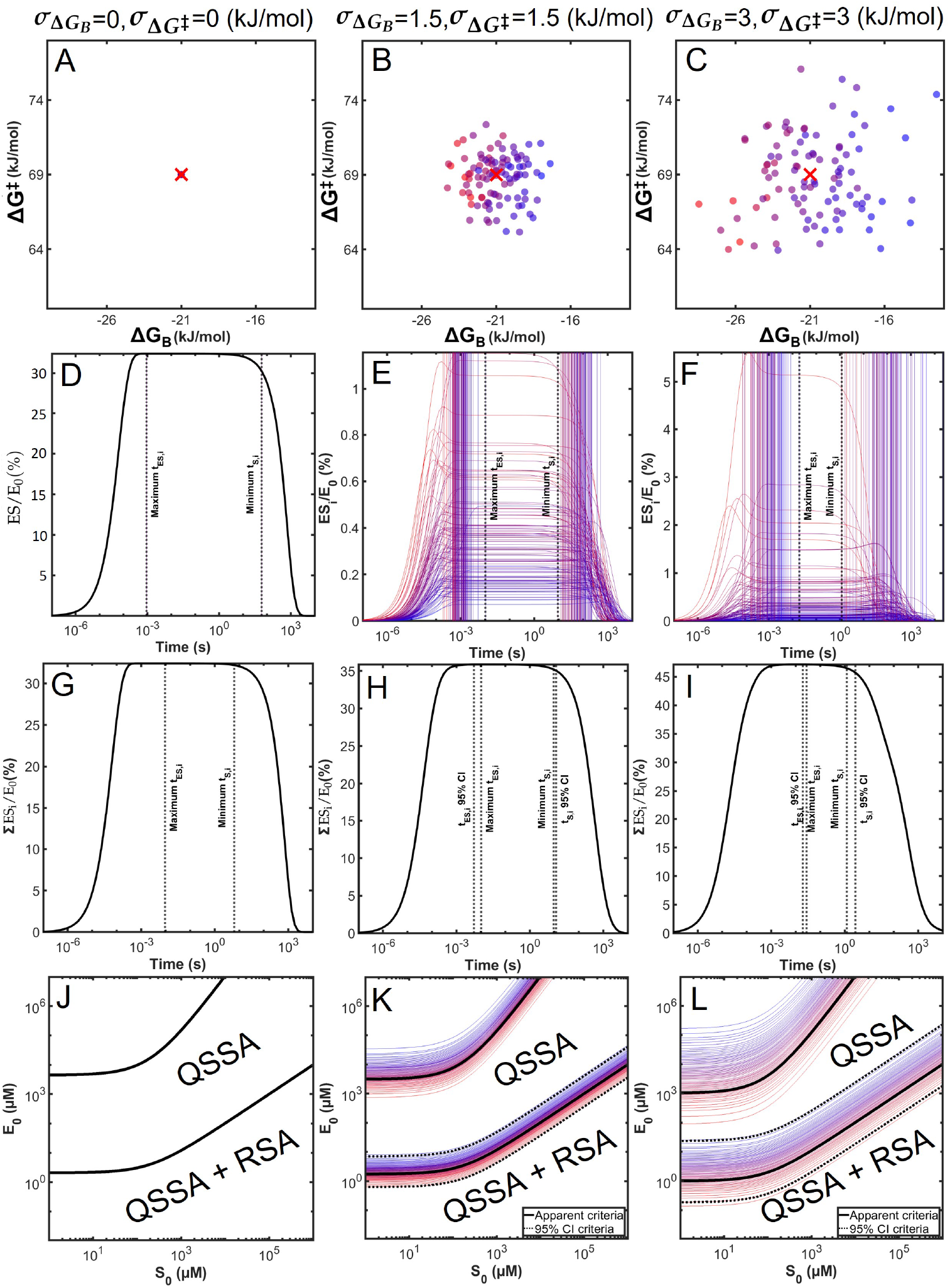
Timescale criteria for QSSA at increasing heterogeneity. Figures A, B and C show a scatterplot of ΔG_B_ and ΔG^‡^. Figures D, E and F show individual concentrations of the different enzyme-substrate complexes ES_i_ over time. The short and long timescale for each substrate are shown with vertical color-scaled lines. The longest of the short timescales (maximum t_ES,i_) and shortest of the long timescales (minimum t_S,i_) are shown with black vertical dashed lines. These dashed lines border the analytical period of steady-state for the enzyme reaction. Figures G, H and I show the total ES concentration, along with the short (t_ES,i_) and long (t_S,i_) timescales as well as 95% CI timescales derived from the distributions. Figures J, K and L show validity plots for the QSSA and RSA criteria for all substrates, using 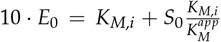. Dashed black lines show the 95% confidence interval for the validity criteria.

For QSSA to hold for all substrates, we must choose a timescale *τ*, such that *t*_*ES,i*_ ≪ *t* ≪ _*S,j*_ for all *i* and *j*. That is, we must choose a timescale such that every enzyme-substrate complex (*ES*_*i*_) has reached its steady-state regime (short timescale), while none have exited it (long timescale), which is fulfilled at time *τ* if *max*(*t*_*ES,i*_) ≪ *τ* ≪ *min*(*t*_*S,i*_). As seen in Figure 8D-I, the timescales given in Eqs. (22) and (23) align with the observed steady-state periods found from the numerical simulations. These observations were used to ensure that a valid timescale was chosen in the numerical simulations in the main article. In practice, this was done by ensuring that all short timescales (Eq. (22)) were at least an order of magnitude below all the long timescales (Eq. (23)), I.E 10*· max*(*t*_*ES,i*_) < *min*(*t*_*S,i*_).

The other assumption used in the derivation of the generalized MM equation was RSA ([*S*_*i*_] ≈*α*_*i*_*S*_0_). Schnell and Mendoza also derived a validity criterion for RSA when multiple substrates are present [23]. Again, using the notation introduced in this article, the validity requirement becomes

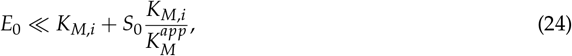

for all substrates *S*_*i*_.

This requirement is more strict than the QSSA criterion, which is illustrated in Figure 8J-L, which show in which regions the upper criteria is an order of magnitude greater than the lower criteria. Thus, if the RSA criteria hold, we can find a valid steady-state timescale for all substrates. To fulfill the criteria given in equation 24, the enzyme concentrations in all the numerical simulations were chosen to be at least an order of magnitude lower than the concentration of all individual substrates (*S*_*i*_). This ensures that the RSA used in the derivation of the generalized MM equation is valid, and that a steady-state period exists.

### Derivation of Apparent MM Parameters from Distributed Energy Landscapes

The apparent MM parameters given by Eqs. (9) and (10) in the main articles become impractical to use as the number of unique substrates increases in the enzyme-catalyzed reaction. However, a simple analytical solution can be derived by letting the substrate heterogeneity be represented by energy distributions. In this representation, heterogeneity manifests itself as a variance in the free energy of the different states along the reaction coordinate in the generalized energy diagram for an enzyme-catalyzed reaction with heterogeneous substrates (see Figure 3 in the main article).

To begin, we will assume that the free energy variations around the states (*G*_*E*+*S*_, *G*_*E*+*S*_‡, *G*_*ES*_, and *G*_*ES*_‡) follow independent probability distributions. We will further assume that the distributions are time-independent, which may happen if either the transitions between the different reaction pathways are fast or slow compared to the time scale of the reaction kinetics. This may be true for substrates that differ only slightly in their energies, such as conformers or substrates with more substantial structural differences, such as isomers, where the transition between the different configurations requires bond breaking.

Using Eqs. (3) and (4) we can rewrite the apparent MM parameters (Eqs. (9) and (10)) in terms of the individual binding (Δ*G*_*B,i*_) and activation 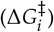 free energy of each substrate.

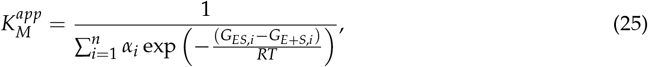

and

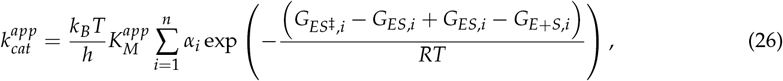

Which reduces to

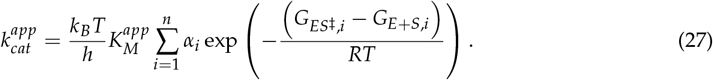

Eqs. (25) and (27) have the same general form: a sum of random variables weighted by their likelihood *α*_*i*_ (molar fraction of substrate *i*). This is the same as the expected value of the variable. 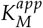 is thus found as

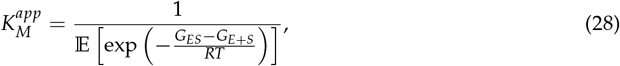

where 𝔼 denotes the expected or mean value of the distribution. We can split the exponential function, using exp (*a* + *b*) = exp (*a*) exp (*b*)

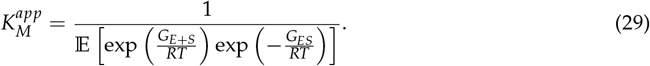

Since we assumed that *G*_*E*+*S*_ and *G*_*ES*_ were independent, the expected value is linear (*E*[*XY*] = *E*[*X*]*E*[*Y*]) and we can rewrite the expression as

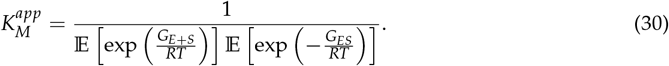

By a similar approach, we find 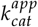 as

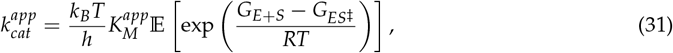

where we again use the properties of the exponential function and the expectation value, to get the following expression

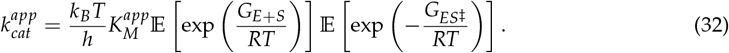

Substituting 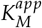 from Eq. (30) gives

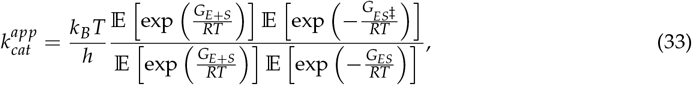

which reduces to

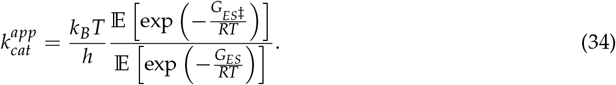

Eqs. (30) and (34) are thus the general solution for the apparent kinetic parameters, 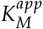 and 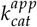^*p*^, for independently distributed energies. We note that these take the form of *E*[exp (*tX*)], which is the Moment Generating Function (MGF) of the random variable X. As such, closed form solutions will exist for any distributions where the MGF exists at 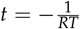 .

### Apparent MM-parameters for normally distributed energy states

We now consider the case where all states are normally distributed. That is 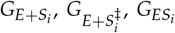, and 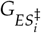 are normal distributed random variable with respective means 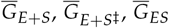, and 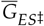 and variances 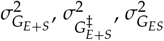 and 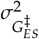. Under these conditions, Eqs. (30) and (34) have analytical solutions.

If the energy of some state, *G*, is normally distributed, then exp 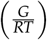 is lognormally distributed. The expected value is then given by: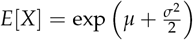. Applying this relationship in the expression for 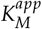 (Eq. (30)) we get

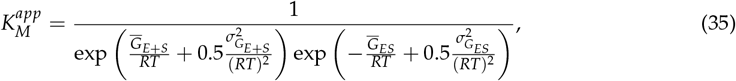

Which we can rewrite as

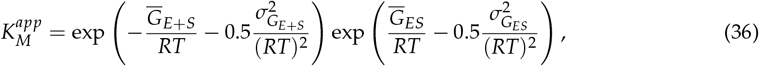

and reduce to

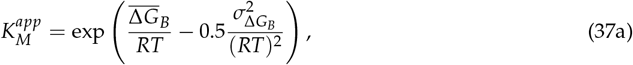

with

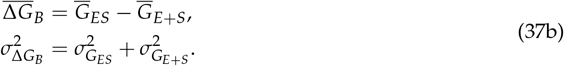

For 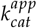 (Eq. (34)) we find

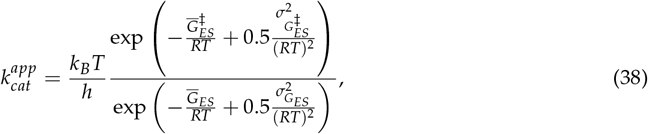

which we can rewrite as

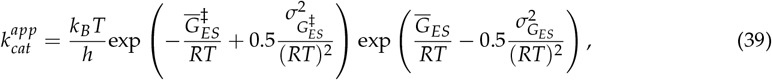

and then reduces to

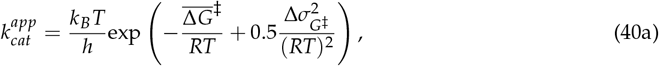

with

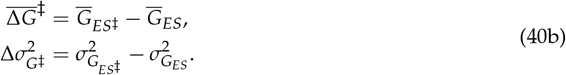

The ratio of Eqs. (40a) and (37a), yields *ϵ*^*app*^

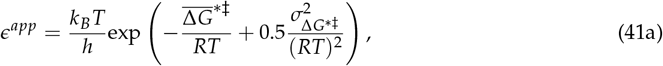

where

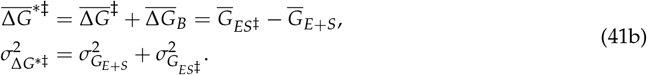

With Eqs. (37a), (40a) and (41a) being the results presented in the article Eqs. (12a), (13a) and (14a).

### Apparent MM-parameters for uniformly distributed energy states

We now consider the case where all states are uniformly distributed. We begin with the simple case of no correlation between Δ*G*_*B*_ and Δ*G*^‡^ (*ρ* = 0). The free energy of the initial state *G*_*E*+*S*_ is uniformly distributed with a range from 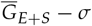 to 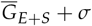, where 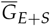 is the mean of the uniform distribution, and *σ* is half of its range. Δ*G*_*B,i*_ will then also be uniformly distributed, with a mean value of 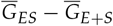 and a range of *σ* as shown in Figure 9.

**Figure 9:**
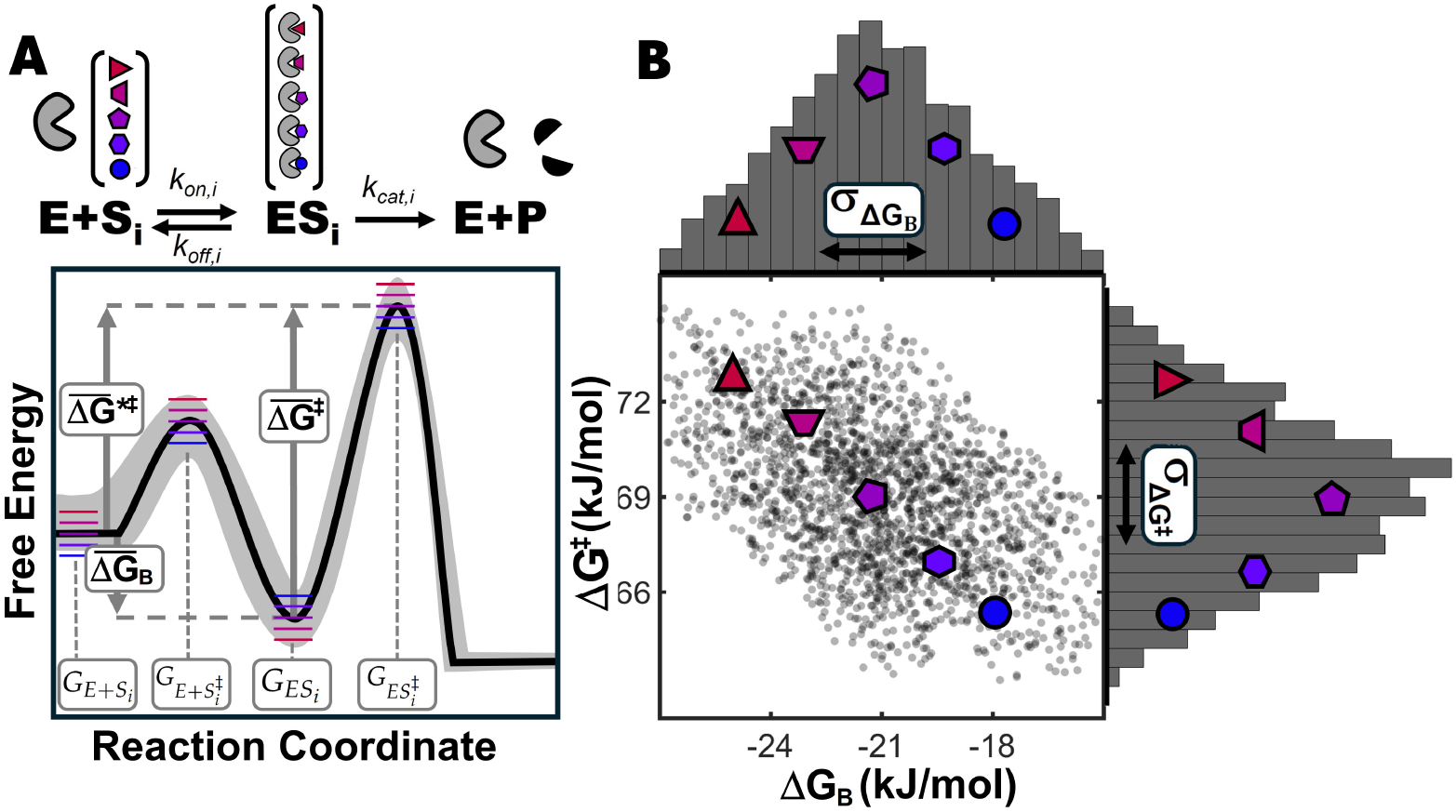
Extension of the classical energy diagram for a homogeneous enzyme-catalyzed reaction to a complex enzyme reaction with multiple non-identical substrates and a shared end product using uniform distributions. A) Heterogeneous substrate energy diagram, where the energy levels of all states are drawn from uniform distributions distributions. B) Scatter plot of the resulting free energy of binding (ΔG_B_) and the free energy of activation (ΔG^‡^) for the energy diagrams. Standard deviations 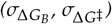 are indicated in the histograms. Note that the two distributions are correlated with a Pearson correlation coefficient of ρ=-0.5, due to the variance of the shared ES_i_ state.

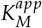 is then given by Eq. (30) as

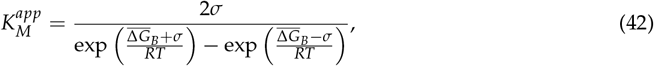

Which reduces to

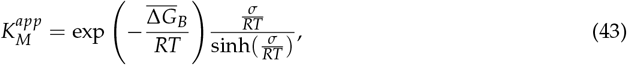

For no correlation 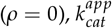 is similarly given by Eq. (34) as

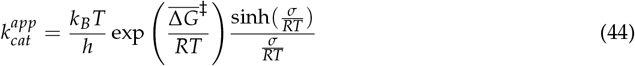

If correlation is introduced 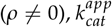 is harder to evaluate continuously, since the convolution of two uniform distributions is not uniformly distributed, except in specific cases.

For correlation of 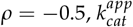 is unaffected by the variance and reduces to

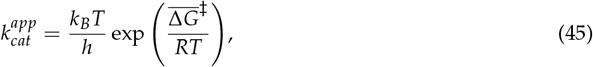

which is analogous to the normally distributed case.

For the complete correlation (*ρ* = −1), the result is

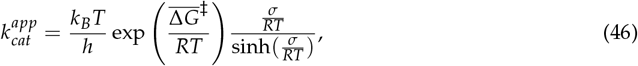

which is identical to Eq. (43), and results in a decrease in 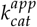 as the variance increases, analogously to the normally distributed case.

In Figure 10 we compared the derived analytical equations for the apparent MM parameters to numerical results, but in this case for uniformly distributed free energies. The numerical simulations were performed using the same methodology as for normal distributions, which is described in the main article.

**Figure 10:**
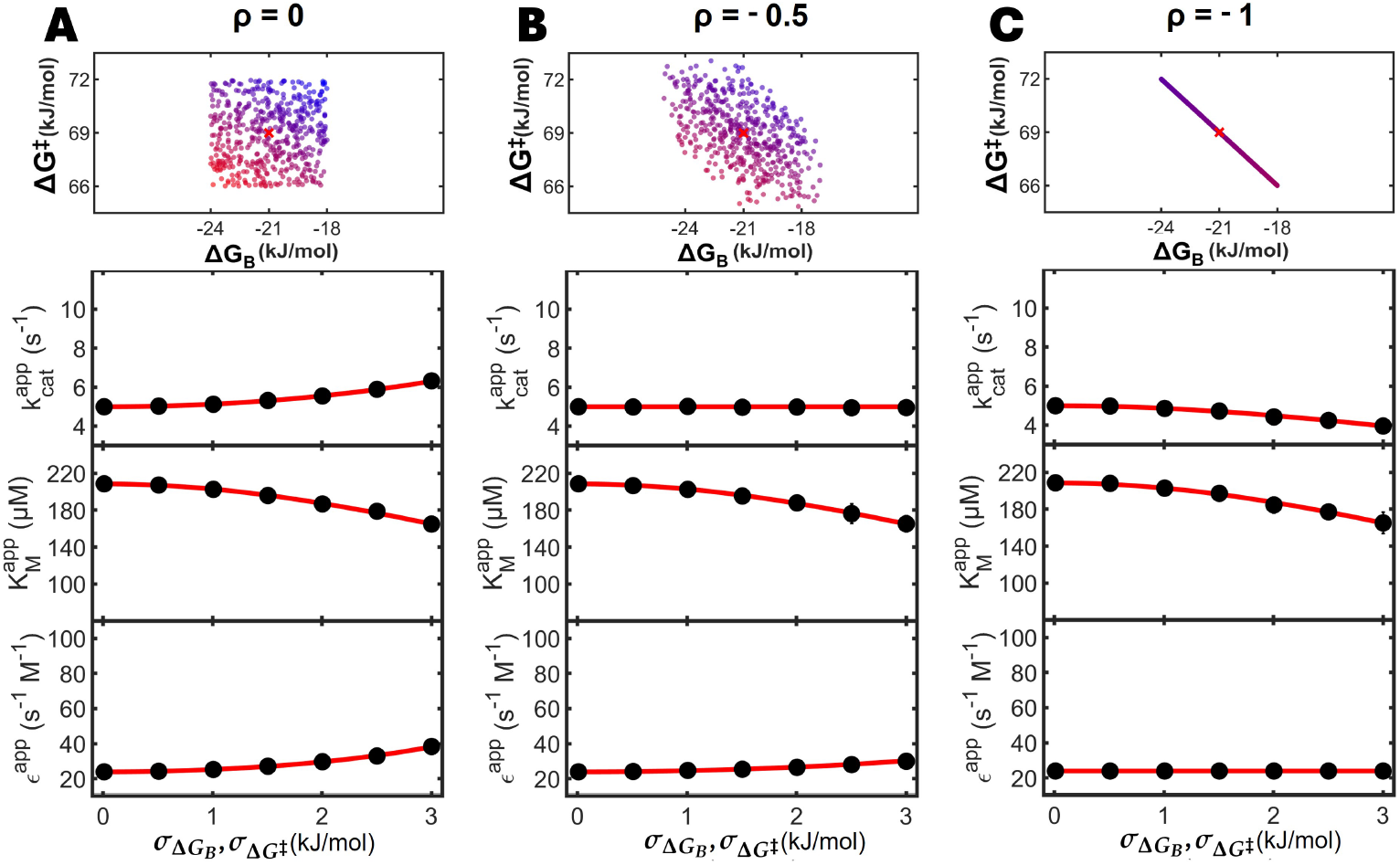
Effect of increasing substrate heterogeneity on the apparent MM parameters, with varying correlations due to variance in the enzyme-substrate complex, using uniformly distributed energies. The kinetic heterogeneity was defined by the standard deviation of the binding free energy (ΔG_B_) and activation free energy (ΔG^‡^). Simulations were performed with fixed mean free energies, 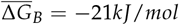 and 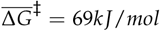, and increasing standard deviation under different levels of correlations. The red curve is the predicted change in the apparent MM parameters using the analytical results of Eqs. 43, 46 and their ratio. The filled circles are estimated apparent MM parameters from numerical simulation with increasing heterogeneity (see Figure 6), and error bars indicate 95% confidence intervals of the simulations. Panel A, B and C illustrate the effect of an increasing correlation between ΔG_B_ and ΔG^‡^, resulting from an increasing proportion of variance in the enzyme-substrate complex. Pearson correlation coefficients between ΔG_B_ and ΔG^‡^ is ρ = 0 (A), ρ = −0.5 (B) and ρ = −1 (C).

### Statistical interpretation of the apparent MM parameters

The results presented in Eqs. (12a), (13a) and (14a) determine how the apparent MM parameters are influenced by the individual substrates, as weighted sums of the individual kinetics parameters. These weighted sums follow the form of different weighted means, which provide insight into the observable kinetics of the mixture, and may provide an easier means of calculating the apparent kinetics of substrate mixtures.

If we in Eq. (9) define the weights as 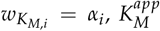 may be seen as a Weighted Harmonic Mean (WHM) of all *K*_*M,i*_ in the substrate mixture

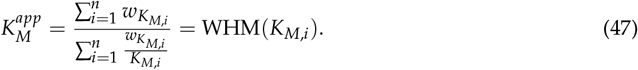

Because the harmonic mean is dominated by the smallest terms, 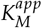 can be substantially smaller than the arithmetic mean of all the *K*_*M,i*_ in the mixture.

Similarly, if we in Eq. (10) define the weights as 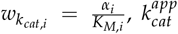 may be viewed as a Weighted sArithmetic Mean (WAM) of all *k*_*cat,i*_ in the substrate mixture

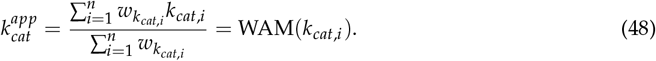

The apparent *k*_*cat*_ value is thus a weighted arithmetic mean of *k*_*cat,i*_ values, with the weight being related to the ratio between a particular substrates abundance and the enzyme affinity of that substrate.

Finally, by combining Eqs. (9) and (10) we find an expression for the apparent specificity constant

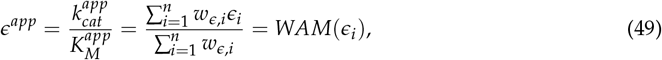

where 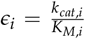 is the specificity constant for substrate *S*_*i*_. As seen from Eq. (49), *ϵ*^*app*^ is a WAM of *ϵ*_*i*_ values with weights *w*_*ϵ,i*_ = *α*_*i*_.

## Notes

### Competing Interest Statement

The authors have declared no competing interest.

